# Ribosome Heterogeneity in Development and Disease

**DOI:** 10.1101/2023.07.25.550527

**Authors:** Yuen Gao, Hongbing Wang

**Author notes:** Correspondence should be addressed to Hongbing Wang at the above address.

## Abstract

The functional ribosome is composed of ∼80 ribosome proteins. With the intensity-based absolute quantification (iBAQ) value, we calculate the stoichiometry ratio of each ribosome protein. We analyze the ribosome ratio-omics (Ribosome^R^), which reflects the holistic signature of ribosome composition, in various biological samples with distinct functions, developmental stages, and pathological outcomes. The Ribosome^R^ reveals significant ribosome heterogeneity among different tissues of fat, spleen, liver, kidney, heart, and skeletal muscles. During tissue development, testes at various stages of spermatogenesis show distinct Ribosome^R^ signatures. During *in vitro* neuronal maturation, the Ribosome^R^ changes reveal functional association with certain molecular aspects of neurodevelopment. Regarding ribosome heterogeneity associated with pathological conditions, the Ribosome^R^ signature of gastric tumors is functionally linked to pathways associated with tumorigenesis. Moreover, the Ribosome^R^ undergoes dynamic changes in macrophages following immune challenges. Taken together, with the examination of a broad spectrum of biological samples, the Ribosome^R^ barcode reveals ribosome heterogeneity and specialization in cell function, development, and disease.

**One-Sentence Summary:** Ratio-omics signature of ribosome deciphers functionally relevant heterogeneity in development and disease.

---

Ribosome is a macromolecular complex to carry out protein synthesis. The eukaryotic ribosome comprises ∼80 highly conserved ribosome proteins (RPs) and 4 rRNAs. It was once considered that ribosome composition is predominantly fixed and homogeneous; the dynamic regulation of protein synthesis during development and pathogenesis is mainly mediated by the activity-dependent alteration of RP-mRNA interaction, post-transcriptional and post-translational modification of mRNA and RP, and epigenetic effects from non-coding RNAs (Gebauer and Hentze, 2004; Song et al., 2021). However, the existence of ribosomes with distinct and different morphology suggests heterogeneity in composition and potentially in function (Norris et al., 2021). Emerging lines of evidence have found that, while some core RPs are central and invariably incorporated into the ribosomes, ribosomes lacking specific RPs are still functional (Shi et al., 2017; Genuth and Barna, 2018). Notably, tissue-specific variations in the expression of particular RPs are observed (Sugihara et al., 2010; Gupta and Warner, 2014; Guimaraes and Zavolan, 2016). Environmental and developmental impacts also affect the level of RP expression and ribosome composition (Reschke et al., 2013; Slavov et al., 2015). These findings suggest that ribosome composition is dynamically regulated and not statically homogeneous (Genuth and Barna, 2018; Barna et al., 2022).

Studies with particular RPs have demonstrated the functional relevance of ribosome heterogeneity. Mutations of specific RPs, including *RPL5, RPL11, RPS17, RPS19, and RPS24*, are associated with Diamond Blackfan Anemia (DBA) (Horos and von Lindern, 2012). It is hypothesized that ribosomes containing distinct RPs may be tailored to preferentially translate a specific pool of mRNAs. Strikingly, knockdown of *Rps19* or *Rpl11* leads to less translation of *Bag1* and *Csde1* mRNA, the protein levels of which are reduced in DBA patient samples (Horos et al., 2012). Beyond translational regulation of individual mRNAs, it is found that specialized ribosomes may be used to translate mRNA pools linked to distinct cellular functions more effectively. For example, the RPS25/eS25- and RPL10A/uL1-containing ribosomes show enrichment of non-overlapping mRNA pools that are associated with functionally different Gene Ontology (GO) groups (Shi et al., 2017).

Considering that previous studies mainly focus on the impact of individual RP on ribosome function, it is unclear whether ribosome heterogeneity only reflects limited outcomes under certain conditions rather than being a general cellular phenomenon. There is little evidence to identify the extensive existence of ribosome heterogeneity. Most importantly, whether and how ribosome heterogeneity is associated with cell identity and the dynamic development and pathogenesis processes is fundamentally unknown. We hypothesize that, due to the complexity of the biological system, functional ribosome heterogeneity involves alteration of many RPs on a genome-wide scale. With a novel bioinformatics approach, we examined the expression ratio of every RP. The ribosome ratio-omics (Ribosome^R^) reveals specific heterogeneity associated with tissue type, *in vivo* and *in vitro* development, and disease. The Ribosome^R^ signature as a genome-wide ribosome barcode identifies a broad extent of ribosome heterogeneity and suggests functional significance.

## MATERIALS AND METHODS

### Datasets

We used the published public proteomic data. The relative expression ratio of RPs in various tissues was analyzed with proteomic data of isolated 80S monosomes from fat, spleen, liver, kidney, heart, and skeletal muscle (Li et al., 2022). The relative expression ratio of RPs in developing testes was analyzed with proteomic data of isolated 80S monosomes from 7-day, 14-day, 28-day, and adult testes (Li et al., 2022). The relative expression ratio of RPs in cultured cortical neurons (Sharma et al., 2015) and human normal and tumor gastric tissues (Ni et al., 2019) was analyzed with whole-cell proteomic data. The relative expression ratio of RPs in control and activated macrophage cells was analyzed with single-cell proteomic data (Woo et al., 2022).

### Ribosome Ratio-omics (Ribosome^R^)

We calculated the expression ratio of each RP to the level of all detected RPs. Depending on the isolation and proteomic detection method, most of the core RPs were detected. Data from various tissues and developing testes reported the expression of 82 and 81 RPs, respectively (Li et al., 2022). Data from cultured cortical neurons (Sharma et al., 2015), human normal and tumor gastric tissues (Ni et al., 2019), and control and activated macrophage cells (Woo et al., 2022) reported the expression of 81, 79, 86, and 71 RPs, respectively. The iBAQ algorithm was used for the ribosomal protein ratio analysis because the iBAQ values are proportional to the molar quantities of the proteins (Woo et al., 2022). The expression ratio of each RP was calculated by (individual RP iBAQ value) / (total RPs iBAQ value). The holistic ribosome ratio-omic (Ribosome^R^) values are reported in Supplemental Table 1-5.

### Principal component analysis (PCA)

The principal component analysis was performed using ClustVis (Metsalu and Vilo, 2015). Unit variance scaling is applied to rows; SVD (singular value decomposition) with imputation is used to calculate principal components. Prediction ellipses are such that, with a probability of 0.95, a new observation from the same group will fall inside the ellipse.

### Correlation analysis

The correlation matrix was calculated in R. Data were plotted using GraphPad Prism 9.

### Heatmap clustering analysis

The Heatmap clustering analysis was performed using ClustVis (Metsalu and Vilo, 2015). Rows are centered; unit variance scaling is applied to rows. Both rows and columns are clustered using correlation distance and average linkage.

### Differential Ribosome^R^ analysis and volcano plot

A significant differential RP ratio was recognized when the adjusted p-value of less than 0.05 was observed. The Volcano plots were made using GraphPad Prism 9.

### Gene ontology (GO) analysis

The GO analysis and enrichment were performed using ShinyGO (Ge et al., 2020). The relevant biological processes were listed in descending order, which was determined by FDR (false discovery rate) and the level of alteration.

### STRING analysis

The STRING database was used to predict potential molecular interactions with the RPs showing ratio alterations. Relevant RPs are placed in the first shell. Likely interacting proteins are placed in the second shell.

### Illustration graphs

Illustration graphs were made with BioRender.

## RESULTS

### Ribosome^R^ reveals tissue-specific ribosome heterogeneity

The tissue-specific function is, in theory, determined by the expression of tissue-specific genes. As the level of mRNA transcripts is not positively correlated with the level of the corresponding proteins, tissue-specific and activity-dependent translation alteration offers necessary control of gene expression. Using the proteomic data of the 80S ribosome (Li et al., 2022), we analyzed the stoichiometry of each core RPs in various tissues. We chose functionally distinct tissues, including fat as an adipose tissue, spleen as an immune tissue, liver and kidney as epithelial tissues, and heart and skeleton muscle as muscle tissues (Fig. 1A). We found that each tissue has a distinct Ribosome^R^, which offers a holistic signature of the stoichiometry of each detected RP (Supplementary Table 1). Principle component analysis (PCA) of Ribosome^R^ identified four functionally distinct groups, namely fat, kidney and liver, heart and skeletal muscle, and spleen (Fig. 1B). An independent correlation analysis showed that fat and spleen are distinctively separated from each other and also from 4 other tissues (Fig. 1C). Moreover, the Ribosome^R^ heatmap revealed 4 prominent clusters with fat, spleen, kidney and liver, and heart and skeletal muscle in each cluster (Fig. 1D). Within the epithelial tissue cluster, kidney and liver are partially separated. Within the muscle cluster, heart and skeletal muscle are partially separated (Fig. 1D). The outcome of our new bioinformatic approach demonstrates that the genome-wide Ribosome^R^ detects tissue-specific ribosome heterogeneity with functional relevance.

**Figure 1.**
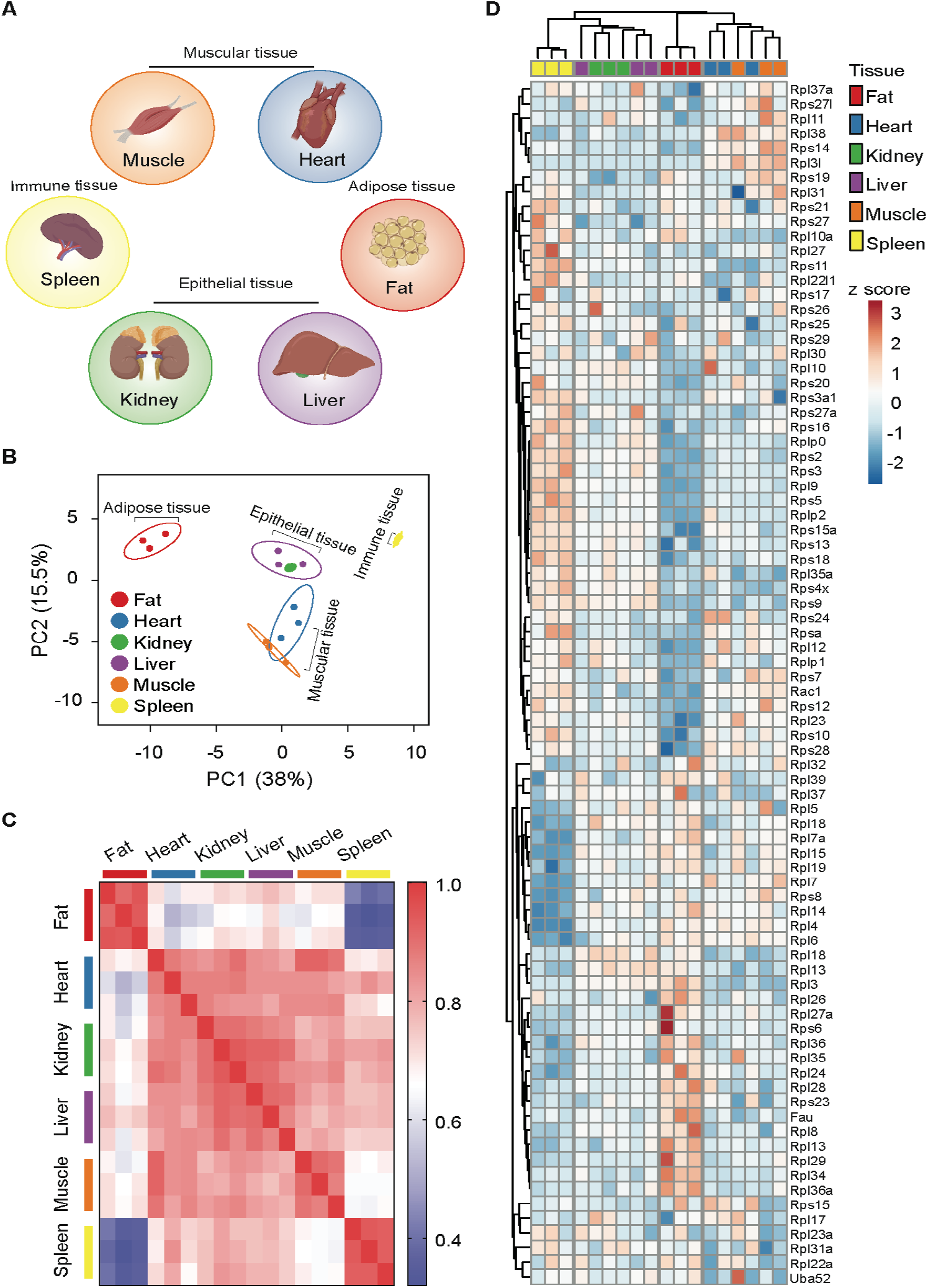
Ribosome^R^ reveals ribosome heterogeneity in tissues with distinct physiological functions. (**A)** Tissues with specialized function. (**B)** Principal component analysis of the expression ratio of each RP in 6 mouse tissues (n=3 for each tissue). The Ribosome^R^ signatures of 82 RPs were compared. **(C)** Correlation matrix of the Ribosome^R^ signatures in 6 mouse tissues. Correlation coefficient of > 0.85 is observed within each tissue. The correlation coefficient between different tissues is expressed as color-coded. **(D)** Heatmap analysis of the Ribosome^R^ signature in 6 mouse tissues. Rows (RP ratio) and columns (distinct tissues) are clustered using correlation distance and average linkage.

We further compared Ribosome^R^ between fat and spleen, which are the most different tissues (Fig. 1B), showing the least correlation (Fig. 1C). Interestingly, the adipose tissue displays a higher expression ratio overwhelmingly in the large subunit RPs (Fig. 2A, 2B, and 2C) and lower expression ratio in the small subunit RPs (Fig. 2A, 2D, and 2E). STRING analysis suggests functional interaction between Mdm2 (murine double minute 2) and Rack1 (receptor for activated C kinase 1) and the RPs showing increased ratio in fat cells (Fig. 2C). Mrpl36, which is a mitochondrial RP, is predicted to interact with RPs showing decreased ratio in fat (Fig. 2E). Relevant to adipose tissue function, Rack1 regulates adipogenesis (Kong et al., 2016) and Mdm2 promotes the onset of fat liver disease (Lin et al., 2022).

**Figure 2.**
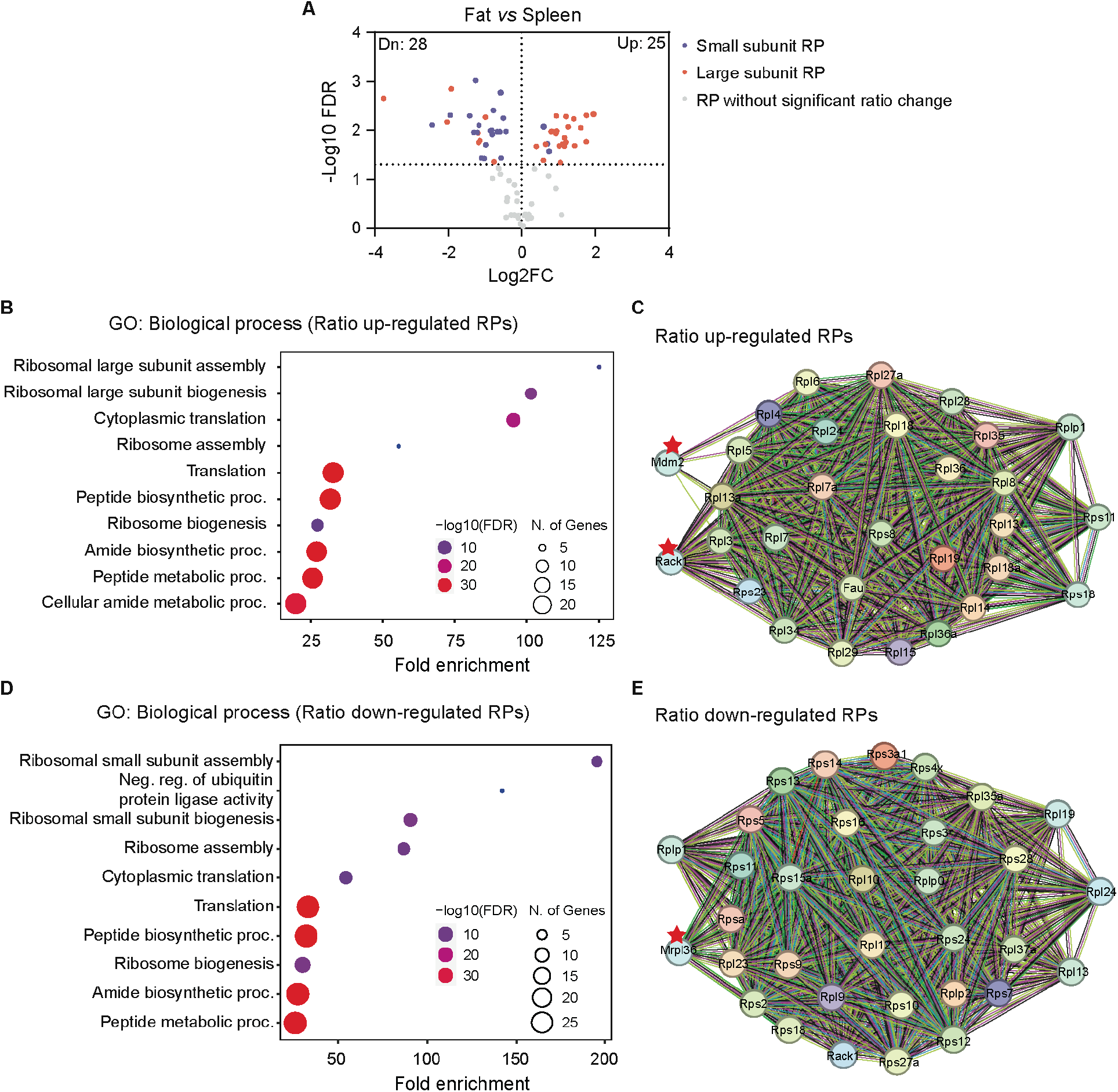
The distinct Ribosome^R^ in fat suggests potential functional relevance to adipogenesis. (**A**) Volcano plot identifies up- and down-regulated expression ratios of specific RPs in fat versus spleen. (**B** and **D**) The top 10 biological processes identified by GO (gene ontology) analysis with the RPs showing up-regulated (**B**) and down-regulated ratios (**D**) in fat versus spleen. (**C** and **E**) STRING analysis identifies the molecular and functional interaction with the RPs that show increased (**C**) and decreased (**E**) stoichiometry in fat versus spleen.

### Ribosome^R^ reveals dynamic ribosome heterogeneity in developing testis

To examine whether there are dynamic changes in ribosome heterogeneity during development, we analyzed the Ribosome^R^ of testes in mice at different postnatal ages (Fig. 3A). With regards to the spermatogenesis process, the 7-day-old testis functionally supports the genesis of spermatogonia, which are 2N cells containing 2 copies of each chromosome. The 14-day and 28-day-old testes functionally support the development of spermatocyte (i.e., 4N cell) and spermatid (i.e., 1N cell), respectively. The adult testis (∼60-day old) contains a mixture of these cells and mature sperm cells. Ribosome^R^ analysis of the proteomic data (Li et al., 2022) found development-specific heterogeneity (Supplementary Table 2). The PCA of Ribosome^R^ separated the testes at different ages into four groups (Fig. 3B). The correlation analysis showed that ribosome stoichiometry in adult testes is in stark contrast to the developing testes at 7, 14, and 28 days of age (Fig. 3C). The Ribosome^R^ heatmap revealed 4 distinct clusters (Fig. 3D). Strikingly, hierarchical clustering indicates that the Ribosome^R^ signature in 7-day testes shows the difference to 14-day, 28-day, and adult testes in ascending order (Fig. 3D). The Ribosome^R^ signature reveals the association of the progressive alteration in ribosome heterogeneity with the progressive and distinctive testis development stages.

**Figure 3.**
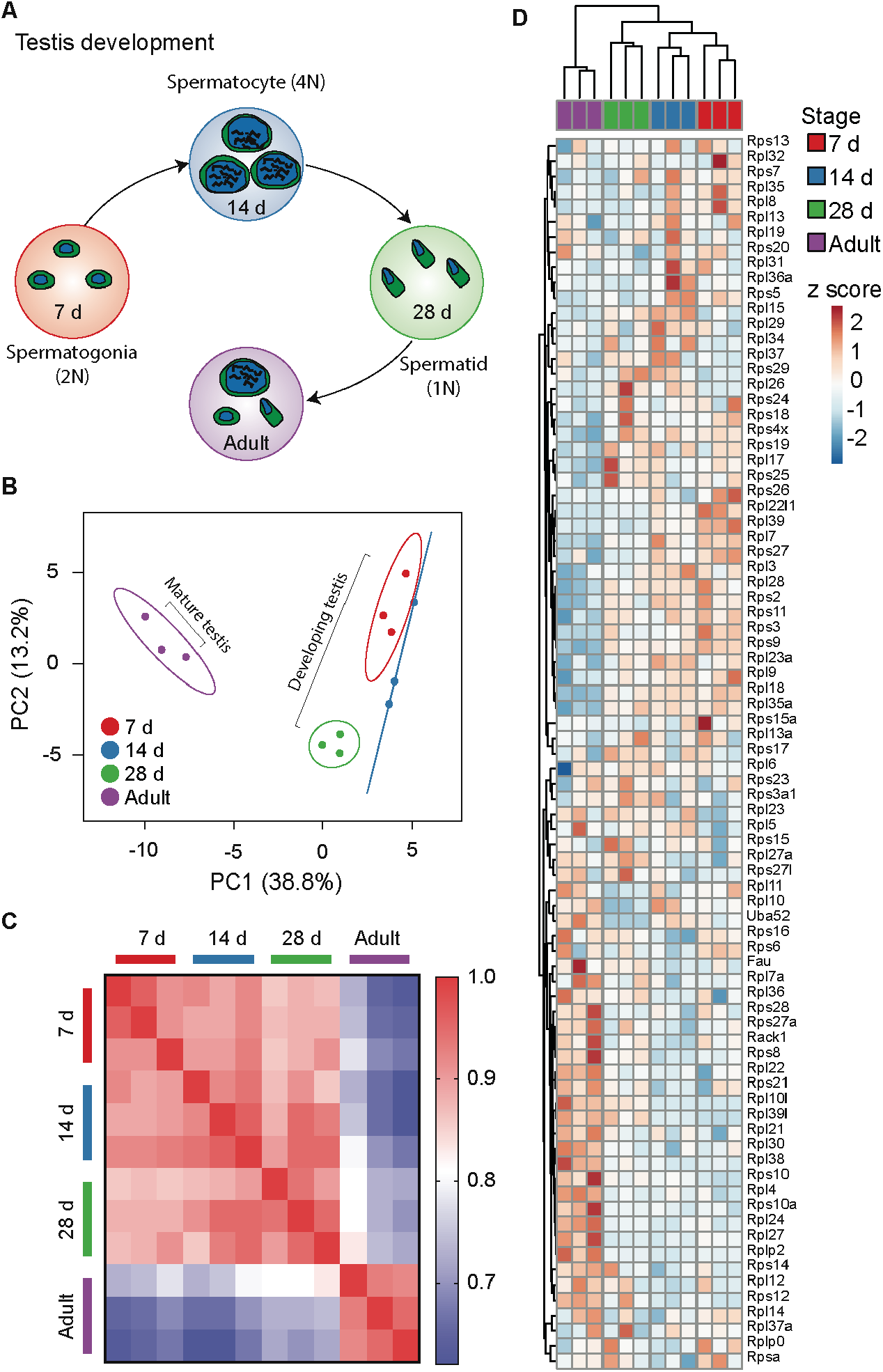
Ribosome^R^ reveals distinct ribosome heterogeneity during testis development. (**A)** Testes at different development stages display distinct cell types during spermatogenesis. (**B)** Principal component analysis of RP ratios in developing and mature mouse testes (n=3 for each group). Ribosome^R^ signatures of 81 RPs were compared. **(C)** Correlation matrix of the Ribosome^R^ signatures in developing and mature mouse testes. Correlation coefficient of > 0.91 is detected within each group. The correlation coefficient between different tissues is expressed as color-coded. (**D)** Heatmap analysis of the Ribosome^R^ signatures in developing and mature mouse testes. Rows are centered; unit variance scaling is applied to rows. Rows (RP ratio) and columns (7-day, 14-day, 28-day, and adult testis) are clustered using correlation distance and average linkage.

### Ribosome^R^ reveals dynamic ribosome heterogeneity during neuronal maturation

To better understand whether there are dynamic changes in ribosome heterogeneity during the development of a specific cell type, we analyzed Ribosome^R^ in cultured cortical neurons (Supplementary Table 3). During the well-characterized *in vitro* maturation, DIV (days in vitro) 5 neurons have moderately developed axons and dendrites with little functional synapses; DIV15 neurons show elaborated axons and dendrites and full-scale synaptogenesis paired with robust expression of synaptic proteins (Fig. 4A).

**Figure 4.**
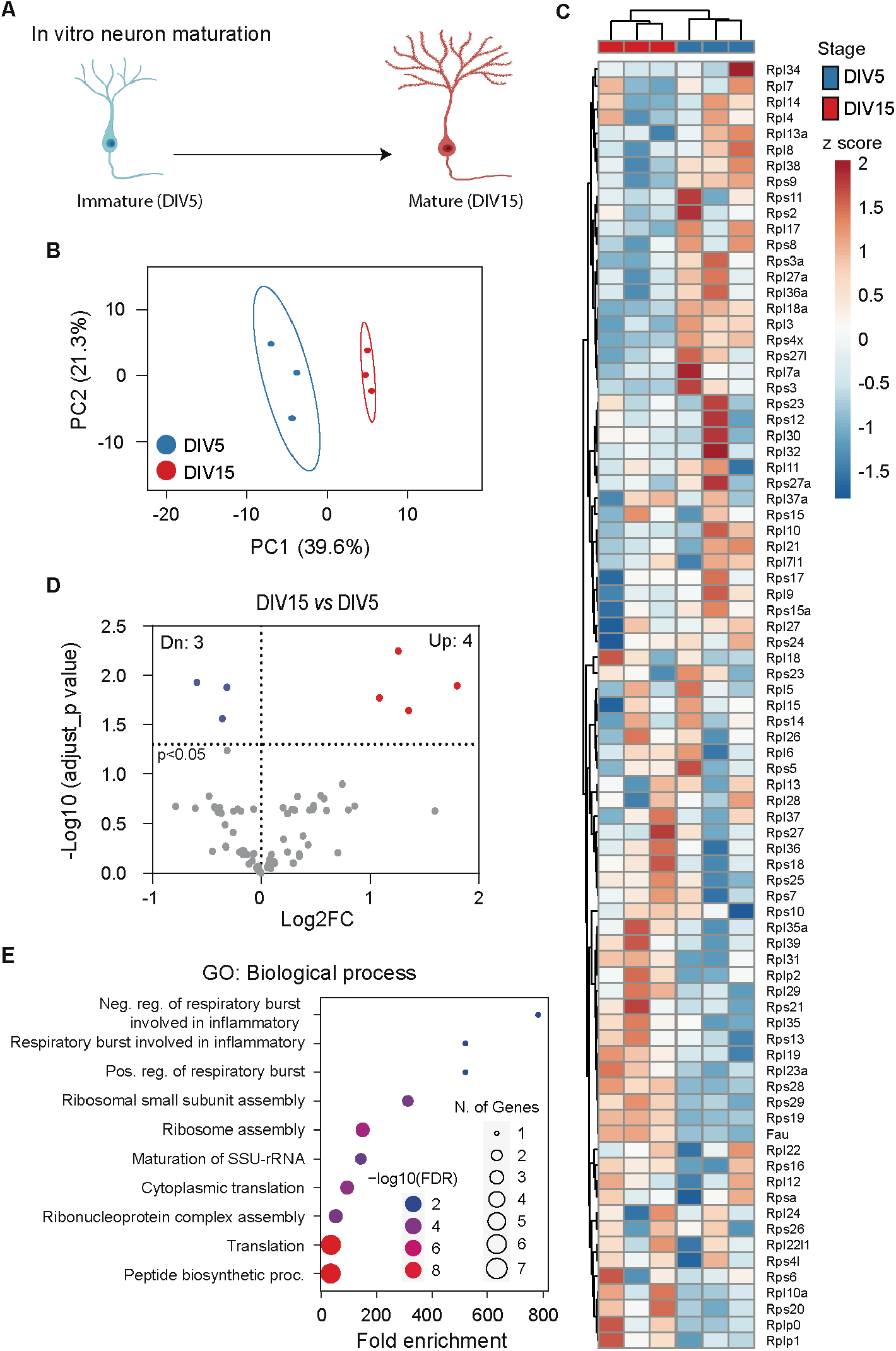
Ribosome^R^ reveals ribosome heterogeneity during *in vitro* neuron maturation. **(A)** Cultured cortical neurons undergo morphological and functional maturation. From DIV (days in vitro) 5 to DIV 15, axons and dendrites become elongated and branched; functional synapses are formed. (**B)** Principal component analysis of RP ratios in DIV 5 and DIV 15 neurons (n=3 for each group). Ribosome^R^ signatures of 81 RPs were compared. **(C)** Heatmap analysis of the Ribosome^R^ signatures in DIV 5 and DIV 15 neurons. Rows (RP ratio) and columns (DIV 5 and DIV 15 neurons) are clustered using correlation distance and average linkage. (**D)** Volcano plot identifies upregulated and down-regulated expression ratios of specific RPs during *in vitro* maturation. (**E)** The top 10 biological processes identified by GO (gene ontology) analysis with the RPs that show ratio alteration during *in vitro* maturation.

We examined the proteomic data collected from DIV 5 and DIV 15 neurons (Sharma et al., 2015). The PCA of Ribosome^R^ showed that the immature DIV 5 neurons and mature DIV 15 neurons display different ribosome compositions (Figure 4B). The heatmap analysis also revealed the Ribosome^R^ signatures in DIV 5 and 15 neurons as distinct clusters (Fig. 4C). By using the differential Ribosome^R^ assay, we found that the mature neurons show increased stoichiometry of 4 RPs and decreased stoichiometry of 3 RPs (Fig. 4D). Go analysis with these 7 RPs revealed that, among the top 10 biological processes, the maturation-induced changes in ribosome heterogeneity are associated with respiratory burst (Fig. 4E). Interestingly, respiratory burst is functional relevant to the release of ROS (reactive oxygen species), the dynamic alterations of which controls synaptic plasticity and neural development (Massaad and Klann, 2011; Oswald et al., 2018a; Oswald et al., 2018b).

### Ribosome^R^ reveals ribosome heterogeneity in tumorigenesis

Compared to normal tissue development, tumorigenesis leads to uncontrolled proliferation and cell immortalization. We examined Ribosome^R^ signatures with proteomic data collected from 82 human normal gastric tissues and 58 human gastric tumor tissues (Fig. 5A) (Ni et al., 2019). The PCA of Ribosome^R^ (Supplementary Table 4) revealed 2 partially separated groups (Fig. 5B). The heatmap analysis revealed 2 main clusters (Fig. 5C). One cluster is enriched with normal gastric tissues (i.e., 76 normal and 10 tumor tissues); another cluster is predominantly consisted of gastric tumor tissues (i.e., 48 tumor and 6 normal tissues) (Fig. 5C). Compared with the normal gastric tissues, the tumor tissues showed increased expression ratio in 34 RPs and decreased ratio in 23 RPs (Fig. 5D). GO analysis with these RPs identified, among the top 10 biological processes, neddylation (Fig. 5E) as a potential link to tumorigenesis (Zhou et al., 2019; Naik et al., 2020). The STRING prediction found that RPs with increased ratio change interact with Fbl (Fibrillarin), Bysl (Bystin or Bystin-Like), and Mdm2 (Fig. 5F), which regulates tumorigenesis through p53 (Alarcon-Vargas and Ronai, 2002; Senturk and Manfredi, 2012). The STRING analysis with the RPs showing decreased ratio suggests functional interaction with Rack1 (Fig. 5G), which regulates various aspects of ribosome function and tumorigenesis (Li and Xie, 2015).

**Figure 5.**
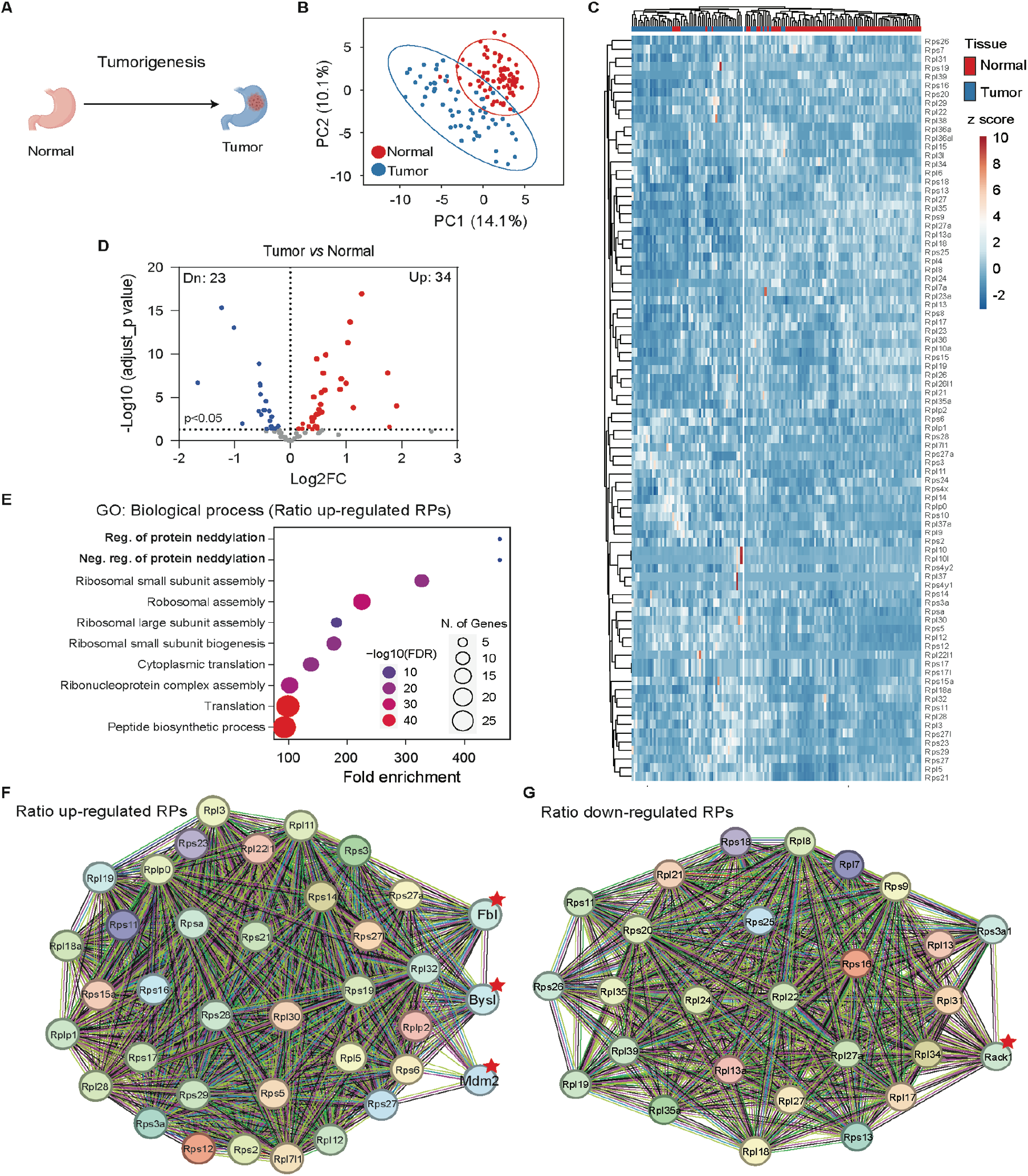
Gastric tissue tumorigenesis shows distinct Ribosome^R^ signatures. (**A)** Tumorigenesis alters cell fate and function in gastric tissue. **(B)** Principal component analysis of RP ratios in human normal gastric tissues (n=82) and gastric tumor tissues (n=58). Proteomic data from normal and tumor human gastric tissues were analyzed. The Ribosome^R^ signature of 86 RPs was compared. (**C)** Heatmap analysis of the Ribosome^R^ signatures in normal and tumor tissues. Rows (RP ratio) and columns (normal and tumor gastric tissues) are clustered using correlation distance and average linkage. (**D)** Volcano plot identifies up- and down-regulated expression ratios of specific RPs in tumor tissue. (**E)** The top 10 biological processes identified by GO (gene ontology) analysis with the RPs that show ratio alteration in the tumor tissues. **(F and G)** STRING analysis and identification of the potential molecular and functional interaction with the RPs that show increased **(F)** and decreased **(G)** stoichiometry in the tumor tissues.

### Ribosome^R^ reveals ribosome heterogeneity in acute macrophage activation

Macrophages, as the early responding immune cells, react to infection or inflammation and neutralize the pathogens (Fig. 6A) (Zhang and Wang, 2014). We analyzed the single-cell proteomic data collected from the murine macrophage cell line RAW 264.7 (Woo et al., 2022). The RAW 264.7 cells were treated with lipopolysaccharide (LPS), a main component of bacteria membrane and a potent reagent to activate macrophage. The single-cell Ribosome^R^ signatures were compared among the 10 control and 38 LPS-challenged cells. The single-cell proteomic data provide reasonable coverage and detect more than 50 RPs in each cell (Supplementary Table 5). The PCA of Ribosome^R^ revealed 2 groups with limited overlap (Fig. 6B). The heatmap analysis revealed 2 main clusters (Fig. 6C). Hierarchical clustering showed that one main cluster contains all control cells in one sub-cluster and a small fraction of the LPS-challenged cells (i.e., 3 cells) in another sub-cluster (Fig. 6C). Another main cluster contains the rest of the LPS-challenged cells (i.e., 35 cells) (Fig. 6C). Compared with the 10 control cells, the 38 LPS-challenged cells showed increased stoichiometry in 9 RPs and decreased stoichiometry in other different 9 RPs (Fig. 6D). The GO analysis showed that, among the top 10 biological processes, the dynamic changes in ribosome heterogeneity are functionally associated with responses to infection (Fig. 6E).

**Figure 6.**
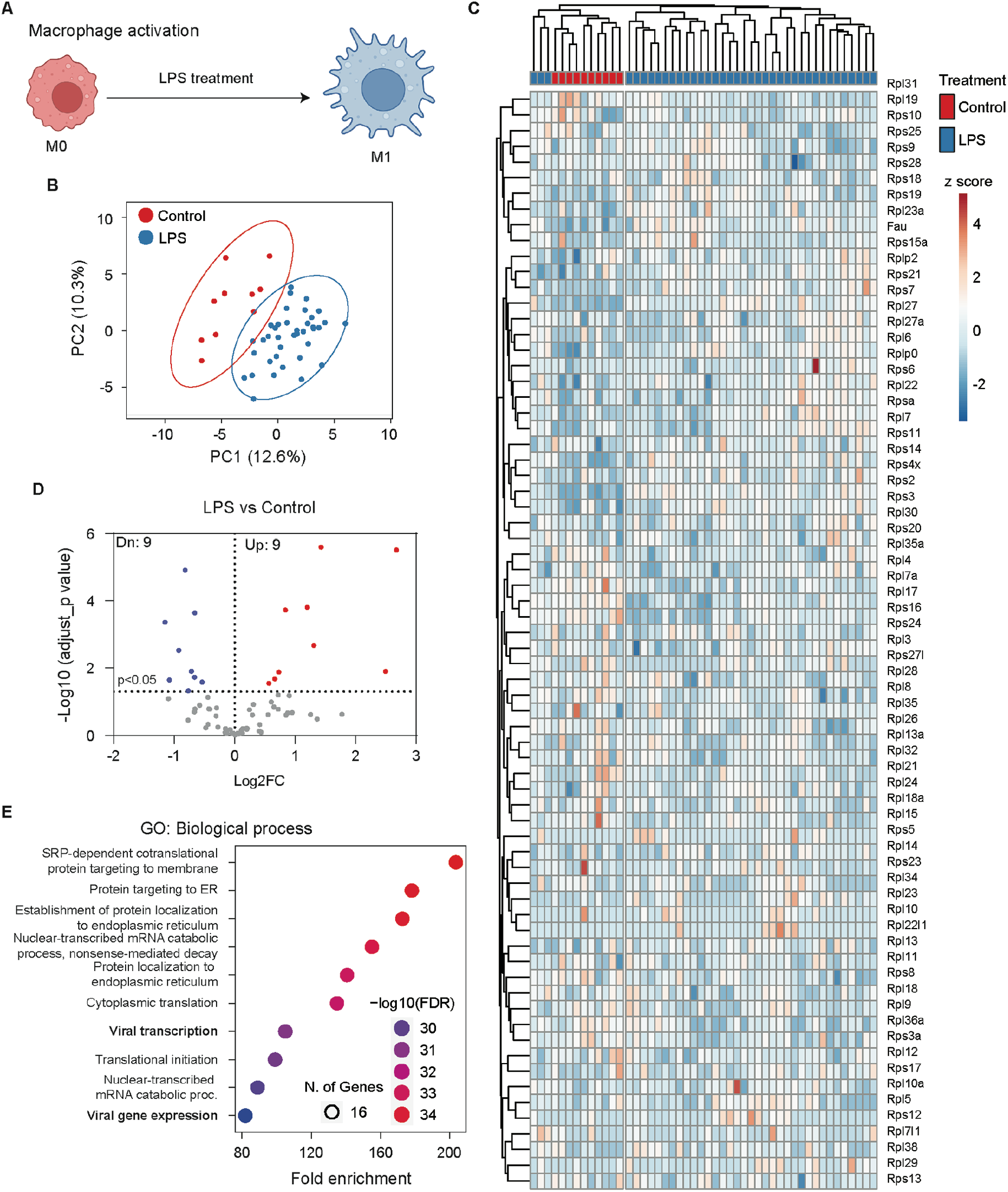
Macrophage cells undergo dynamic Ribosome^R^ changes following activation. **(A)** The proinflammatory activator LPS (lipopolysaccharide) stimulates macrophage cells, leading to immune responses such as the secretion of cytokines. **(B)** Principal component analysis of RP ratios in control (n=10) and LPS-treated macrophages (n=38). The Ribosome^R^ signature of 71 RPs was compared. **(C)** Heatmap analysis of Ribosome^R^ signatures in control and LPS-treated macrophage cells. Rows (RP ratio) and columns (control and LPS-treated cells) are clustered using correlation distance and average linkage. **(D)** Volcano plot identifies upregulated and down-regulated expression ratio of specific RPs in the LPS-treated cells. (**E)** The top 10 biological processes identified by GO (gene ontology) analysis with the RPs that show ratio alteration in the LPS-treated cells.

## DISCUSSION

Precise and dynamic regulation of gene expression is essential for cell function and development. Given that variation of the specific RPs affects mRNA translation and is associated with pathology and disease, the functional relevance of ribosome heterogeneity has been proposed but is yet under debate. Here, we performed the first genome-wide and holistic characterization of ribosome composition in various tissues under physiological and pathological conditions. With Ribosome^R^, we identified ribosome heterogeneity associated with tissue type and development. We also detected alteration of Ribosome^R^ in disease conditions. Our findings implicate a broad existence of ribosome heterogeneity and its functional relevance. We also expect that Ribosome^R^, as an effective analysis method, will be useful to decipher an essential and new dimension of gene expression regulation.

Previous studies have observed differential expression of specific RP mRNAs in various mammalian tissues (Bortoluzzi et al., 2001; Gupta and Warner, 2014; Guimaraes and Zavolan, 2016). Notably, the mRNA levels of about 25% of the RP genes show tissue-specific expression (Guimaraes and Zavolan, 2016) but also see (Gupta and Warner, 2014). The functional relevance of the mRNA heterogeneity is not clear. The mRNA expression profile does not render separated PCA groups with distinct tissue functions. The hierarchical clustering analysis shows that the functionally similar tissues are not in the same sub-cluster but are somewhat distant. For example, the RP mRNA signature in kidney (an epithelial tissue) is more similar to that in adipose tissue than liver (another epithelial tissue) (Guimaraes and Zavolan, 2016). Sugihara et al. analyzed the ribosome proteomic data collected from liver, testis, and mammary gland. They found differential levels of several RP-like proteins but failed to detect tissue-specific ribosome heterogeneity (Sugihara et al., 2010). Here, we use Ribosome^R^ and, for the first time, identify functionally relevant and tissue-specific heterogeneity. The adipose, epithelial, muscular, and immune tissue show distinct Ribosome^R^. We notice that Ribosome^R^ is insufficient to detect significant heterogeneity among tissues of a similar type. For example, heart and skeletal muscle, as well as liver and kidney, show slightly overlapping Ribosome^R^. It is unclear whether the mixed cell types within each tissue lead to compromised resolution of Ribosome^R^. When comparing Ribosome^R^ in 2 tissues with obviously different physiological functions, we identified striking differences in ribosome composition between fat and spleen. The Ribosome^R^ analysis further suggests functional relevance of the tissue-specific heterogeneity, as indicated by the role of Rack1 and Mdm2 in adipogenesis and fat liver disease.

If ribosome heterogeneity is associated with biological function, the ribosome composition in a specific tissue is expected to dynamically cope with the functional changes during development. Even with the testis tissue consisting of different cell types, Ribosome^R^ detects the interaction of development and function in 4 distinct development stages. For the cultured primary cells consisting of mainly neurons rather than mixed cell types, Ribosome^R^ detects a difference between the immature and mature neurons. The maturation-associated ribosome changes detect GO functions related to ROS production. While ROS affects neural development and maturation (Wilson et al., 2018), ROS also shows excessive production in DBA (Diamond-Blackfan anemia) (Rio et al., 2019). Interestingly, among the 4 RPs showing an increased ratio in the mature DIV 15 neurons, RPS19, RPS28, and RPS29 mutations have been found in DBA (Horos and von Lindern, 2012; Dokal et al., 2022). Causally, deficient RPS19, of which the mutations are most frequent in DBA (Draptchinskaia et al., 1999), directly results in anemia in mice (Debnath et al., 2017).

Ribosome composition and function alteration has been observed in cancer and tumorigenesis (Guimaraes and Zavolan, 2016; Kang et al., 2021). Targeting ribosomes is also a potential cancer therapy (Gilles et al., 2020). Guimaraes and Zavolan analyzed The Cancer Genome Atlas (TCGA) data and found that certain RP genes, such as the *RPL39L*, are commonly dysregulated in different cancer types; whether cancer cells and normal cells show distinct ribosome heterogeneity remains unclear (Guimaraes and Zavolan, 2016). Our Ribosome^R^ analysis with a large proteomic dataset consisting of 82 normal and 58 gastric tumor tissues revealed 2 separate clusters enriched with normal and tumor samples, respectively. Among the 34 RPs showing an increased ratio in cancer, RPL11 and RPL5 missense mutations occur in 73% and 66% of the 19000 cancer samples across 49 cancer types (Orsolic et al., 2020). While an increase in RPS15A associates with the progression of various cancers (Guo et al., 2018), a decrease in RPS15A inhibits cancer cell proliferation (Zhao et al., 2015; Yao et al., 2016). Among the 23 RPs showing decreased ratio in cancer, RPL22 is reduced through heterozygous deletion in ∼ 10% T-ALL (T-acute lymphoblastic leukemia) (Rao et al., 2012). Various mutations in RPL22 are also found in solid cancers (Ferreira et al., 2014). While it is not clear whether the tumor-specific ribosome heterogeneity regulates the translation of tumor-specific proteins or reflects a pathological outcome, STRING prediction of the RPs with increased ratio functional association with the Mdm2/p53 axis, Fbl, and Bysl. Among the Ribosome^R^-identified tumor-associated RPs with increased ratio, RPL5, RPL11, RPS3, RPS14, RPS19, RPS27, RPS27A, and RPS27L have been found to physically interact with Mdm2 (Kang et al., 2021), which inhibits the function of the tumor suppressor p53 (Alarcon-Vargas and Ronai, 2002; Senturk and Manfredi, 2012). Beyond its well-known role in regulating transcription, the Mdm2/p53 axis, which affects and responds to nucleolus activity (Liu et al., 2016), may also participate in the ribosome assembly process. The Ribosome^R^-identified RP-Fbl interaction may suggest altered ribosome biogenesis in gastric cancer (Nguyen Van Long et al., 2022). Bysl is involved in rRNA processing and 40S ribosome biogenesis during development and cancer cell proliferation (Adachi et al., 2007; Wang et al., 2009). Rack1, which is predicted by STRING to functionally interact with the RPs showing decreased ratio, stably associates with ribosomes (Johnson et al., 2019) and regulates the translation of specific mRNA pools (Majzoub et al., 2014; Thompson et al., 2016). Intriguingly, RACK1 affects cancer progression in a tissue-dependent manner. While it promotes breast and lung cancers (Li and Xie, 2015), it suppresses gastric tumors (Deng et al., 2012).

Single-cell Ribosome^R^ analysis reveals dynamic ribosome alteration in macrophages following immune challenge. Interestingly, the Ribosome^R^ in each cell does not respond equally to LPS. The 3 LPS-challenged cells showed different Ribosome^R^ compared to both the control cells and the other 35 LPS-challenged cells. This finding is consistent with the emerging observation that macrophages are epigenetically diversified (Ma et al., 2022). The different single-cell Ribosome^R^ signatures in LPS-stimulated macrophages further suggest functional diversity.

In summary, Ribosome^R^ identifies the broad existence of ribosome heterogeneity with functional relevance to development and disease. The Ribosome^R^ offers a new strategy to analyze ribosome composition and function.

## ACKNOWLEDGMENTS

This research was supported by NIH grants R01MH124992 and R01MH119149 (to HW).

## SUPPLEMENTARY TABLE LEGENDS

**Supplementary Table 1**. Expression ratio of RPs in different mouse tissues of fat, heart, skeletal muscle, kidney, liver, and spleen.

**Supplementary Table 2**. Expression ratio of RPs in mouse testes at different ages.

**Supplementary Table 3**. Expression ratio of RPs in DIV5 and DIV15 neurons.

**Supplementary Table 4**. Expression ratio of RPs in human normal and tumor gastric tissues.

**Supplementary Table 5**. Single-cell expression ratio of RPs in control and LsPS-treated RAW 264.7 cells.

## REFERENCES

Adachi, K., Soeta-Saneyoshi, C., Sagara, H., and Iwakura, Y. (2007). Crucial role of Bysl in mammalian preimplantation development as an integral factor for 40S ribosome biogenesis. Mol Cell Biol 27(6), 2202–2214. doi: 10.1128/MCB.01908-06.

Alarcon-Vargas, D., and Ronai, Z. (2002). p53-Mdm2--the affair that never ends. Carcinogenesis 23(4), 541–547. doi: 10.1093/carcin/23.4.541.

Barna, M., Karbstein, K., Tollervey, D., Ruggero, D., Brar, G., Greer, E.L., et al. (2022). The promises and pitfalls of specialized ribosomes. Mol Cell 82(12), 2179–2184. doi: 10.1016/j.molcel.2022.05.035.

Bortoluzzi, S., d’Alessi, F., Romualdi, C., and Danieli, G.A. (2001). Differential expression of genes coding for ribosomal proteins in different human tissues. Bioinformatics 17(12), 1152–1157. doi: 10.1093/bioinformatics/17.12.1152.

Debnath, S., Jaako, P., Siva, K., Rothe, M., Chen, J., Dahl, M., et al. (2017). Lentiviral Vectors with Cellular Promoters Correct Anemia and Lethal Bone Marrow Failure in a Mouse Model for Diamond-Blackfan Anemia. Mol Ther 25(8), 1805–1814. doi: 10.1016/j.ymthe.2017.04.002.

Deng, Y.Z., Yao, F., Li, J.J., Mao, Z.F., Hu, P.T., Long, L.Y., et al. (2012). RACK1 suppresses gastric tumorigenesis by stabilizing the beta-catenin destruction complex. Gastroenterology 142(4), 812–823 e815. doi: 10.1053/j.gastro.2011.12.046.

Dokal, I., Tummala, H., and Vulliamy, T. (2022). Inherited bone marrow failure in the pediatric patient. Blood 140(6), 556–570. doi: 10.1182/blood.2020006481.

Draptchinskaia, N., Gustavsson, P., Andersson, B., Pettersson, M., Willig, T.N., Dianzani, I., et al. (1999). The gene encoding ribosomal protein S19 is mutated in Diamond-Blackfan anaemia. Nat Genet 21(2), 169–175. doi: 10.1038/5951.

Ferreira, A.M., Tuominen, I., van Dijk-Bos, K., Sanjabi, B., van der Sluis, T., van der Zee, A.G., et al. (2014). High frequency of RPL22 mutations in microsatellite-unstable colorectal and endometrial tumors. Hum Mutat 35(12), 1442–1445. doi: 10.1002/humu.22686.

Ge, S.X., Jung, D., and Yao, R. (2020). ShinyGO: a graphical gene-set enrichment tool for animals and plants. Bioinformatics 36(8), 2628–2629. doi: 10.1093/bioinformatics/btz931.

Gebauer, F., and Hentze, M.W. (2004). Molecular mechanisms of translational control. Nat Rev Mol Cell Biol 5(10), 827–835. doi: 10.1038/nrm1488.

Genuth, N.R., and Barna, M. (2018). The Discovery of Ribosome Heterogeneity and Its Implications for Gene Regulation and Organismal Life. Mol Cell 71(3), 364–374. doi: 10.1016/j.molcel.2018.07.018.

Gilles, A., Frechin, L., Natchiar, K., Biondani, G., Loeffelholz, O.V., Holvec, S., et al. (2020). Targeting the Human 80S Ribosome in Cancer: From Structure to Function and Drug Design for Innovative Adjuvant Therapeutic Strategies. Cells 9(3). doi: 10.3390/cells9030629.

Guimaraes, J.C., and Zavolan, M. (2016). Patterns of ribosomal protein expression specify normal and malignant human cells. Genome Biol 17(1), 236. doi: 10.1186/s13059-016-1104-z.

Guo, P., Wang, Y., Dai, C., Tao, C., Wu, F., Xie, X., et al. (2018). Ribosomal protein S15a promotes tumor angiogenesis via enhancing Wnt/beta-catenin-induced FGF18 expression in hepatocellular carcinoma. Oncogene 37(9), 1220–1236. doi: 10.1038/s41388-017-0017-y.

Gupta, V., and Warner, J.R. (2014). Ribosome-omics of the human ribosome. RNA 20(7), 1004–1013. doi: 10.1261/rna.043653.113.

Horos, R., Ijspeert, H., Pospisilova, D., Sendtner, R., Andrieu-Soler, C., Taskesen, E., et al. (2012). Ribosomal deficiencies in Diamond-Blackfan anemia impair translation of transcripts essential for differentiation of murine and human erythroblasts. Blood 119(1), 262–272. doi: 10.1182/blood-2011-06-358200.

Horos, R., and von Lindern, M. (2012). Molecular mechanisms of pathology and treatment in Diamond Blackfan Anaemia. Br J Haematol 159(5), 514–527. doi: 10.1111/bjh.12058.

Johnson, A.G., Lapointe, C.P., Wang, J., Corsepius, N.C., Choi, J., Fuchs, G., et al. (2019). RACK1 on and off the ribosome. RNA 25(7), 881–895. doi: 10.1261/rna.071217.119.

Kang, J., Brajanovski, N., Chan, K.T., Xuan, J., Pearson, R.B., and Sanij, E. (2021). Ribosomal proteins and human diseases: molecular mechanisms and targeted therapy. Signal Transduct Target Ther 6(1), 323. doi: 10.1038/s41392-021-00728-8.

Kong, Q., Gao, L., Niu, Y., Gongpan, P., Xu, Y., Li, Y., et al. (2016). RACK1 is required for adipogenesis. Am J Physiol Cell Physiol 311(5), C831–C836. doi: 10.1152/ajpcell.00224.2016.

Li, H., Huo, Y., He, X., Yao, L., Zhang, H., Cui, Y., et al. (2022). A male germ-cell-specific ribosome controls male fertility. Nature 612(7941), 725–731. doi: 10.1038/s41586-022-05508-0.

Li, J.J., and Xie, D. (2015). RACK1, a versatile hub in cancer. Oncogene 34(15), 1890–1898. doi: 10.1038/onc.2014.127.

Lin, H., Wang, L., Liu, Z., Long, K., Kong, M., Ye, D., et al. (2022). Hepatic MDM2 Causes Metabolic Associated Fatty Liver Disease by Blocking Triglyceride-VLDL Secretion via ApoB Degradation. Adv Sci (Weinh) 9(20), e2200742. doi: 10.1002/advs.202200742.

Liu, Y., Deisenroth, C., and Zhang, Y. (2016). RP-MDM2-p53 Pathway: Linking Ribosomal Biogenesis and Tumor Surveillance. Trends Cancer 2(4), 191–204. doi: 10.1016/j.trecan.2016.03.002.

Ma, R.Y., Black, A., and Qian, B.Z. (2022). Macrophage diversity in cancer revisited in the era of single-cell omics. Trends Immunol 43(7), 546–563. doi: 10.1016/j.it.2022.04.008.

Majzoub, K., Hafirassou, M.L., Meignin, C., Goto, A., Marzi, S., Fedorova, A., et al. (2014). RACK1 controls IRES-mediated translation of viruses. Cell 159(5), 1086–1095. doi: 10.1016/j.cell.2014.10.041.

Massaad, C.A., and Klann, E. (2011). Reactive oxygen species in the regulation of synaptic plasticity and memory. Antioxid Redox Signal 14(10), 2013–2054. doi: 10.1089/ars.2010.3208.

Metsalu, T., and Vilo, J. (2015). ClustVis: a web tool for visualizing clustering of multivariate data using Principal Component Analysis and heatmap. Nucleic Acids Res 43(W1), W566–570. doi: 10.1093/nar/gkv468.

Naik, S.K., Lam, E.W., Parija, M., Prakash, S., Jiramongkol, Y., Adhya, A.K., et al. (2020). NEDDylation negatively regulates ERRbeta expression to promote breast cancer tumorigenesis and progression. Cell Death Dis 11(8), 703. doi: 10.1038/s41419-020-02838-7.

Nguyen Van Long, F., Lardy-Cleaud, A., Carene, D., Rossoni, C., Catez, F., Rollet, P., et al. (2022). Low level of Fibrillarin, a ribosome biogenesis factor, is a new independent marker of poor outcome in breast cancer. BMC Cancer 22(1), 526. doi: 10.1186/s12885-022-09552-x.

Ni, X., Tan, Z., Ding, C., Zhang, C., Song, L., Yang, S., et al. (2019). A region-resolved mucosa proteome of the human stomach. Nat Commun 10(1), 39. doi: 10.1038/s41467-018-07960-x.

Norris, K., Hopes, T., and Aspden, J.L. (2021). Ribosome heterogeneity and specialization in development. Wiley Interdiscip Rev RNA 12(4), e1644. doi: 10.1002/wrna.1644.

Orsolic, I., Bursac, S., Jurada, D., Drmic Hofman, I., Dembic, Z., Bartek, J., et al. (2020). Cancer-associated mutations in the ribosomal protein L5 gene dysregulate the HDM2/p53-mediated ribosome biogenesis checkpoint. Oncogene 39(17), 3443–3457. doi: 10.1038/s41388-020-1231-6.

Oswald, M.C., Brooks, P.S., Zwart, M.F., Mukherjee, A., West, R.J., Giachello, C.N., et al. (2018a). Reactive oxygen species regulate activity-dependent neuronal plasticity in Drosophila. Elife 7. doi: 10.7554/eLife.39393.

Oswald, M.C.W., Garnham, N., Sweeney, S.T., and Landgraf, M. (2018b). Regulation of neuronal development and function by ROS. FEBS Lett 592(5), 679–691. doi: 10.1002/1873-3468.12972.

Rao, S., Lee, S.Y., Gutierrez, A., Perrigoue, J., Thapa, R.J., Tu, Z., et al. (2012). Inactivation of ribosomal protein L22 promotes transformation by induction of the stemness factor, Lin28B. Blood 120(18), 3764–3773. doi: 10.1182/blood-2012-03-415349.

Reschke, M., Clohessy, J.G., Seitzer, N., Goldstein, D.P., Breitkopf, S.B., Schmolze, D.B., et al. (2013). Characterization and analysis of the composition and dynamics of the mammalian riboproteome. Cell Rep 4(6), 1276–1287. doi: 10.1016/j.celrep.2013.08.014.

Rio, S., Gastou, M., Karboul, N., Derman, R., Suriyun, T., Manceau, H., et al. (2019). Regulation of globin-heme balance in Diamond-Blackfan anemia by HSP70/GATA1. Blood 133(12), 1358–1370. doi: 10.1182/blood-2018-09-875674.

Senturk, E., and Manfredi, J.J. (2012). Mdm2 and tumorigenesis: evolving theories and unsolved mysteries. Genes Cancer 3(3-4), 192–198. doi: 10.1177/1947601912457368.

Sharma, K., Schmitt, S., Bergner, C.G., Tyanova, S., Kannaiyan, N., Manrique-Hoyos, N., et al. (2015). Cell type- and brain region-resolved mouse brain proteome. Nat Neurosci 18(12), 1819–1831. doi: 10.1038/nn.4160.

Shi, Z., Fujii, K., Kovary, K.M., Genuth, N.R., Rost, H.L., Teruel, M.N., et al. (2017). Heterogeneous Ribosomes Preferentially Translate Distinct Subpools of mRNAs Genome-wide. Mol Cell 67(1), 71–83 e77. doi: 10.1016/j.molcel.2017.05.021.

Slavov, N., Semrau, S., Airoldi, E., Budnik, B., and van Oudenaarden, A. (2015). Differential Stoichiometry among Core Ribosomal Proteins. Cell Rep 13(5), 865–873. doi: 10.1016/j.celrep.2015.09.056.

Song, P., Yang, F., Jin, H., and Wang, X. (2021). The regulation of protein translation and its implications for cancer. Signal Transduct Target Ther 6(1), 68. doi: 10.1038/s41392-020-00444-9.

Sugihara, Y., Honda, H., Iida, T., Morinaga, T., Hino, S., Okajima, T., et al. (2010). Proteomic analysis of rodent ribosomes revealed heterogeneity including ribosomal proteins L10-like, L22-like 1, and L39-like. J Proteome Res 9(3), 1351–1366. doi: 10.1021/pr9008964.

Thompson, M.K., Rojas-Duran, M.F., Gangaramani, P., and Gilbert, W.V. (2016). The ribosomal protein Asc1/RACK1 is required for efficient translation of short mRNAs. Elife 5. doi: 10.7554/eLife.11154.

Wang, H., Xiao, W., Zhou, Q., Chen, Y., Yang, S., Sheng, J., et al. (2009). Bystin-like protein is upregulated in hepatocellular carcinoma and required for nucleologenesis in cancer cell proliferation. Cell Res 19(10), 1150–1164. doi: 10.1038/cr.2009.99.

Wilson, C., Munoz-Palma, E., and Gonzalez-Billault, C. (2018). From birth to death: A role for reactive oxygen species in neuronal development. Semin Cell Dev Biol 80, 43–49. doi: 10.1016/j.semcdb.2017.09.012.

Woo, J., Clair, G.C., Williams, S.M., Feng, S., Tsai, C.F., Moore, R.J., et al. (2022). Three-dimensional feature matching improves coverage for single-cell proteomics based on ion mobility filtering. Cell Syst 13(5), 426–434 e424. doi: 10.1016/j.cels.2022.02.003.

Yao, Y., Liu, Y., Lv, X., Dong, B., Wang, F., Li, J., et al. (2016). Down-regulation of ribosomal protein S15A inhibits proliferation of human glioblastoma cells in vivo and in vitro via AKT pathway. Tumour Biol 37(4), 4979–4990. doi: 10.1007/s13277-015-4323-0.

Zhang, L., and Wang, C.C. (2014). Inflammatory response of macrophages in infection. Hepatobiliary Pancreat Dis Int 13(2), 138–152. doi: 10.1016/s1499-3872(14)60024-2.

Zhao, X., Shen, L., Feng, Y., Yu, H., Wu, X., Chang, J., et al. (2015). Decreased expression of RPS15A suppresses proliferation of lung cancer cells. Tumour Biol 36(9), 6733–6740. doi: 10.1007/s13277-015-3371-9.

Zhou, L., Jiang, Y., Luo, Q., Li, L., and Jia, L. (2019). Neddylation: a novel modulator of the tumor microenvironment. Mol Cancer 18(1), 77. doi: 10.1186/s12943-019-0979-1.

